# Phase Separation and Ageing of Glycine-Rich Protein from Tick Adhesive

**DOI:** 10.1101/2023.03.27.534361

**Authors:** Ketan A. Ganar, Polina Turbina, Manali Nandy, Chang Chen, Dennis Suylen, Stan van der Beelen, Emily Louise Pascoe, Constantianus J.M. Koenraadt, Ingrid Dijkgraaf, Siddharth Deshpande

**Author notes:** shared authorship.

## Abstract

Hard ticks feed on their host for multiple days. To ensure firm attachment, they secrete a protein-rich saliva that eventually forms a solid cement cone. The underlying mechanism of this liquid-to-solid transition is not yet understood. This study focuses on the phase transitions of a disordered glycine-rich protein (GRP) that is prominent in tick saliva. We show that GRP undergoes liquid-liquid phase separation via simple coacervation to form biomolecular condensates in salty environments. Cation-pi and pi-pi interactions near the C-terminus promote coacervation while a negatively charged N-terminus prolongs its onset through electrostatic repulsion. Interestingly, GRP condensates exhibit ageing and undergo liquid-to-gel transition to form viscoelastic networks as well as solid-like condensates. Lastly, we provide evidence for protein-rich condensates in natural tick saliva. Our findings provide a starting point to gain insights into the bioadhesion of ticks, develop novel tick control strategies, and towards biomedical applications such as tissue sealants.

## Introduction

Biological adhesives are sticky materials used by a wide variety of organisms for different purposes such as attachment, prey capture, locomotion, building, and defense^1^. The range of species that can produce bioadhesives is diverse, from bacterial biofilms and slimy garden slugs to stickleback nests and sticky spiderwebs^1^. Many animals use protein-based adhesives: sandcastle worms build reef-like mounds^2^, mussels attach via proteinaceous threads^3^, and velvet worms eject adhesive slime to entangle their prey^4^. While some bioadhesives have been studied in considerable detail, knowledge on adhesive mechanisms of many others is significantly lacking. One of the unexplored and unique bioadhesives is produced by ticks, a widespread parasite of public health and economic importance^5, 6^.

Ticks are arthropods that feed on their host by sucking blood over a prolonged period, usually multiple days in the case of hard ticks^7^. Importantly, the prolonged contact and transfer of saliva from the tick to the host can lead to pathogen transmission, which can result in infectious diseases in both humans and animals worldwide, notable examples being Lyme borreliosis in humans, and babesiosis, anaplasmosis, and heartwater in bovine species^8, 9^. To feed successfully, ticks first need to securely attach to the host skin, which they do so in two stages: an initial mechanical attachment, followed by bioadhesive production known as the cement cone^10^ (**Figure 1a**). The mechanical attachment is achieved using the mouthparts which consist of two palps that perform sensory functions, a pair of extendable chelicerae to cut into the host tissue, and a hypostome that acts as a channel for blood as well as to penetrate the outermost layer of the epidermis^10, 11^ (**Supplementary Figure 1**). Once the hypostome has passed through the first skin layer, saliva is secreted from the salivary glands. This milky-white proteinaceous fluid has adhesive properties and undergoes a liquid-to-solid transition once exposed to air, therefore bearing the name cement^10, 12, 13^. A cement cone, resembling a wedge-shaped anchor, is formed around the incision site of the tick which strengthens attachment to the host and facilitates long feeding periods^10, 12, 13^. Adhesive production is commonly associated with hard ticks, belonging to the family *Ixodidae*, allowing them to remain attached to their host for up to 10 days. Thereafter, feeding begins with the formation of small blood pools, followed by intermittent ejection of saliva^10, 14^. For hydration purposes, tick saliva is also highly hygroscopic and salt-rich which helps to absorb moisture from air^14^. However, the mechanism as well as the identity of the key salivary components that are responsible for the formation of such a strong bioadhesive remains unknown.

**Figure 1:**
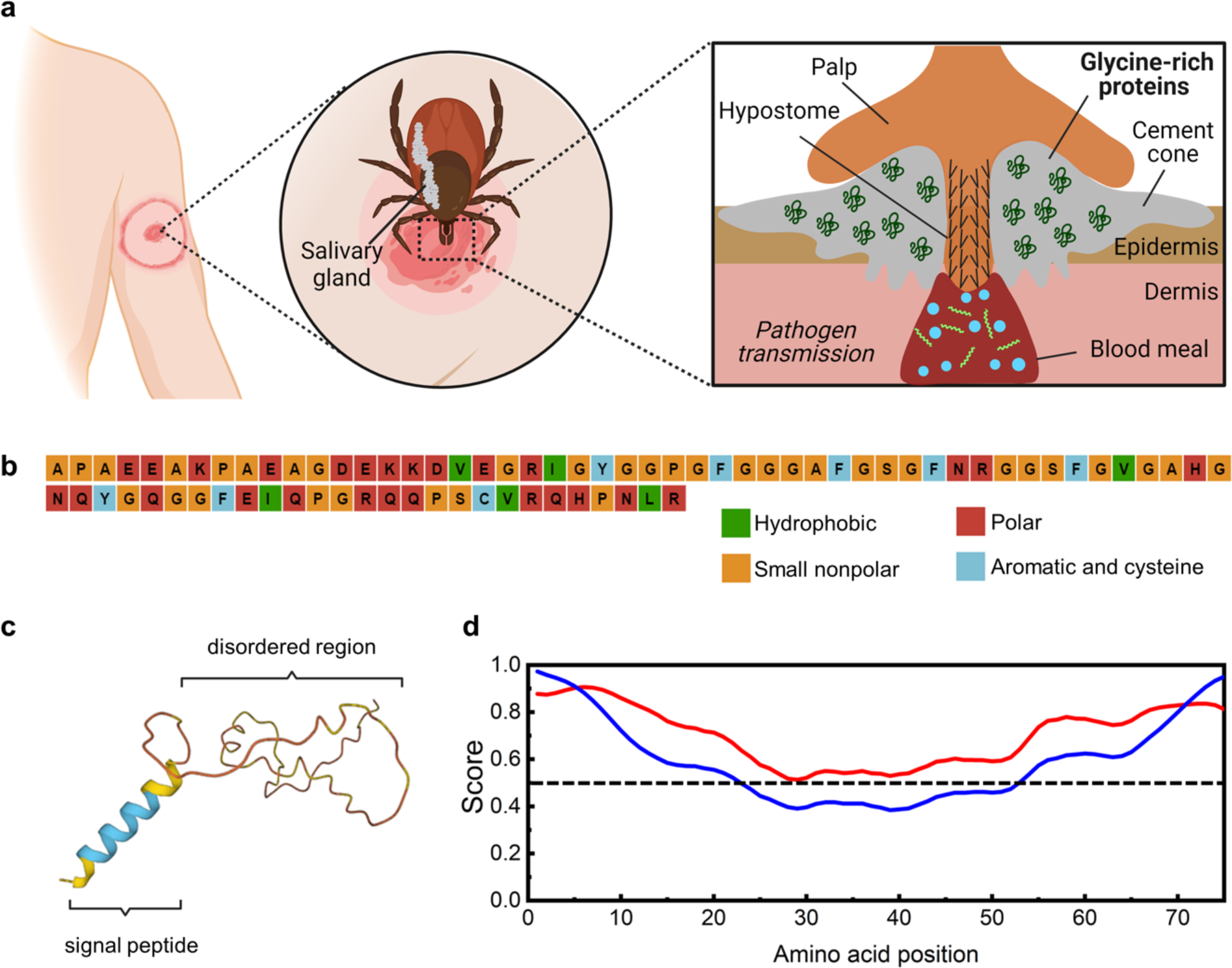
Glycine-rich protein (GRP) present in the tick saliva is intrinsically disordered and shows a high propensity for LLPS. **(a)** Schematic representation showing the consequence of a tick bite. The tick inserts its hypostome into the host epidermis and secretes a protein-rich saliva, abundant in glycine-rich proteins (GRPs). The saliva undergoes liquid-to-solid transition forming a hard cement cone, allowing the tick to feed on the host over several days and facilitating pathogen transmission (the shown “bull’s eye” rash is typical in case of Borrelia infection). **(b)** Amino acid composition of tick-GRP77 shows a high proportion of non-polar (44%, out of which 26% are glycines) and polar amino acids (36%). (**c**) AlphaFold correctly predicts the N-terminal signal peptide of GRP as an α-helix, while the rest of the sequence (tick-GRP77) remains unstructured, indicating a disordered region. (**d**) IUPred3 long disorder mode (red line) scores the entire tick-GRP77 sequence above 0.5, while short disorder score (blue line) shows prominent disorder near the termini, overall indicating tick-GRP77 as a highly disordered protein.

Biochemical and bioinformatic analyses have revealed that glycine-rich proteins (GRPs) are abundant in tick saliva^15, 16, 17, 18^. GRPs have been associated with various physiological functions including providing strength, insolubility and stabilization to the cement cone^10^, but the mechanism by which GRPs facilitate these functions remains unknown. We noted that glycine-rich regions are commonly present in intrinsically disordered proteins (IDPs) since they can prevent protein folding due to their small size and high degree of freedom^19^. Since IDPs do not have a native state, they usually lack secondary structures and can therefore undergo large-scale conformational changes^20^. IDPs have been associated with liquid-liquid phase separation (LLPS) and the formation of biomolecular condensates because of their capacity to establish multiple interactions with neighbouring molecules due to their multivalency^21^. LLPS manifested via coacervation is driven by the interactions between multivalent biomolecules such as proteins and nucleic acids, resulting in polymer-rich droplets that are in equilibrium with a polymer-depleted phase^22^. Coacervates can be simple (single component) or complex (multicomponent) and have been found to play crucial roles in diverse cellular functions^23, 24, 25, 26, 26^. While usually liquid-like, coacervates have been shown to undergo liquid-to-solid transition^4, 27, 28, 29, 30^. This strong link between IDPs, LLPS, and liquid-to-solid transition prompted our attention to study the possible role of tick GRPs in cement cone formation.

Recently, LLPS has indeed been shown to play a role in bioadhesion via liquid-to-solid transition^1^. Various aquatic and terrestrial organisms like mussels, sandcastle worm, spiders, and velvet worms have been shown to utilize LLPS to produce strong adhesives under specific environmental triggers such as cross-linking, pH changes, and evaporation^2, 4, 29, 31, 32, 33, 34, 35^. Once deposited into the host skin, tick saliva is also subjected to various biological triggers: increased local concentrations of salt and salivary proteins, likely aided by evaporation-induced water loss during the secretion, together with a significant temperature rise and a pH drop^36^. Interestingly, all these triggers are known to induce LLPS of proteins^22, 24, 28^.

In this study, we have provided the first experimental evidence that a tick GRP present in the saliva of *Ixodes scapularis* (Uniprot: Q4PME3) undergoes LLPS in the form of simple coacervation and forms GRP-rich condensates. Instead of protein extraction from the cement cone, which is very difficult due to the insolubility of cement cones and the harsh conditions required to isolate the proteins^1, 37^, we use solid-phase peptide synthesis (SPPS) and native chemical ligation (NCL) to synthesize GRP, allowing us to investigate its biochemical properties under controlled *in vitro* conditions. Through structure prediction analysis, we show that GRP is a predominantly disordered protein and has a high propensity of undergoing LLPS. Using a combination of fluorescence microscopy and microfluidics, we demonstrate that GRP undergoes evaporation- and salt-induced LLPS, resulting in simple coacervates with a characteristic fluid-like behaviour. We further show that these condensates age over time to form solid-like aggregates and viscoelastic networks. Finally, we show evidence for the presence of protein-rich condensates in natural saliva from the closely related *Ixodes ricinus* tick. In conclusion, our work illustrates the ability of tick GRP to undergo liquid-to-solid transition through LLPS. These findings can shed further light on the poorly studied cement cone formation and may partly explain the role of GRPs in the saliva in the form of providing necessary adhesive properties for a successful attachment to the host. The obtained knowledge may become relevant for exploring chemicals that hinder tick attachment, consequently aiding the prevention and management of tick-associated diseases. Considering the close interactions of the cement cone with biological tissues, the findings may also prove useful for biotechnological applications, such as medical sealants.

## Results and Discussion

### Sequence and structure prediction analysis suggests GRP to be an intrinsically disordered protein

We selected one of the prominent GRPs present in tick saliva (**Figure 1a**), simply known as GRP (Uniprot: Q4PME3) and refer to it as such here onwards. GRP is a relatively short protein containing 96 amino acid residues. The first 19 N-terminal amino acids (sequence: ^1^MNRMFVLAATLALVGMVFA^19^) constitute the signal peptide necessary for its translocation. The remaining 77 amino acids constitute the mature GRP sequence (20–96) and we refer to it as tick-GRP77 here onwards (to simplify things, amino acid locations within tick-GRP77 are renumbered, i.e., 1^st^ position refers to the 20^th^ amino acid from GRP and 77^th^ is the 96^th^ one). As the name suggests, tick-GRP77 is rich in glycine residues (≈26%) with most of them located in the middle of the sequence (13–63). Further sequence composition analysis by PSIPRED^38^ revealed a large fraction of other non-polar (≈44%; mainly alanine and proline) and polar (≈36%; mainly glutamic acid, arginine, and glutamine) amino acids. In comparison, the fraction of hydrophobic (≈7%; valine and isoleucine) and aromatic (≈9%; tyrosine and phenylalanine) residues was relatively small (**Figure 1b**). We also checked the GRP structure via Alphafold^39^, an algorithm capable of computationally predicting protein structures. Indeed, the GRP sequence showed two distinct regions (**Figure 1c**): The N-terminus of GRP, corresponding to the signal peptide, shows an α-helical conformation. However, the remaining sequence (corresponding to tick-GRP77) gave a very low prediction score, hinting at a lack of any kind of secondary structure. IUPred3 algorithm^40^, which predicts IDRs, also determined the entire tick-GRP77 to be completely disordered using long disorder mode (score above 0.5 throughout the sequence) while the short disorder mode predicted prominent disorder at both the termini (**Figure 1d**). We further evaluated the sequence using CIDER^41^, which plots the fraction of negatively charged versus positively charged residues. Based on this, proteins can be grouped into five different regions according to the diagram of states **(Supplementary Figure 2**). The tick-GRP77 sequence falls in the region of weak polyampholytes and polyelectrolytes, similar to many well-characterized phase-separating IDR-containing proteins like FUS^42^ and viral nucleocapsid^43^. Cumulatively, our bioinformatic analysis indicated a strong inclination for tick-GRP77 to undergo LLPS.

### Tick-GRP77 undergoes liquid-liquid phase separation via simple coacervation

Upon confirming the disordered nature of tick-GRP77 via structure prediction analysis, we proceeded with gathering experimental evidence for its LLPS behavior. We synthesized tick-GRP77 via SPPS and NCL (see Materials and Methods for details). Since the secretion of tick saliva locally enriches its components in the host tissue, aided by water loss through evaporation, we mimicked this via a straightforward droplet evaporation assay (**Figure 2a**). We deposited a small sessile droplet (2 μL) of buffered tick-GRP77 solution (16-500 μM) on a glass slide and let it evaporate at ambient temperature. The non-uniform evaporation of the droplet led to what is commonly known as the coffee-ring effect^44^, concentrating tick-GRP77 at the droplet boundary. Briefly, higher evaporation rate at the air-water-glass triple contact point (the droplet boundary) results in fluid flow towards the boundary to replenish the depleting liquid, bringing along the tick-GRP77 molecules, increasing their concentration at the boundary. For better visualization, we added 5 mol% fluorescently labelled tick-GRP, OG488-GRP77 (fluorescent label: Oregon Green 488; see Materials and Methods for details), to the sample. **Figure 2b** shows a time-lapse, near the droplet boundary, of a 32 μM tick-GRP77 solution in phosphate buffer saline (PBS; 137 mM NaCl, 2.7 mM KCl, 8 mM Na_2_HPO_4_, and 2 mM KH_2_PO_4_, pH 7.4) evaporating over time, while **Figure 2c** shows the same time-lapse in fluorescence (**Supplementary Movie 1**). The droplet initially spread over the hydrophilic glass slide and after approximately 30 seconds, an outer boundary was pinned due to the rough glass surface.

**Figure 2:**
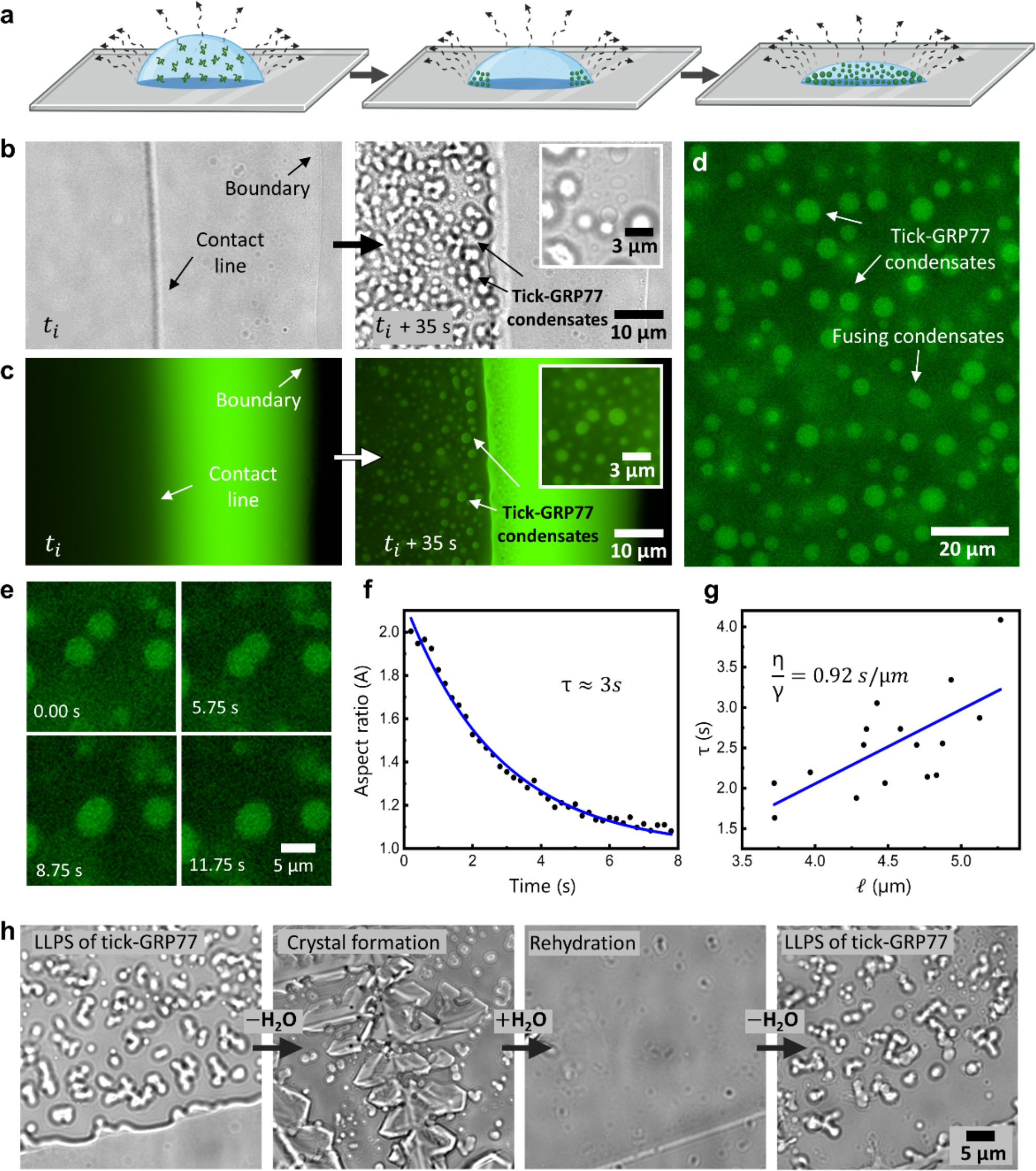
Tick-GRP77 undergoes liquid-liquid phase separation via simple coacervation to form liquid-like condensates. **(a)** Schematic of the droplet evaporation assay, where a droplet of a buffered tick-GRP77 solution is allowed to evaporate at room temperature, continuously increasing the protein concentration at the droplet boundary. (**b-c**) Evaporation of a 2 μL tick-GRP77 (32 μM; prepared in PBS, pH 7.4) droplet leads to the formation of spherical condensates near the contact line (*t*_*i*_ = 15 min) and the condensate formation rapidly spreads inward. Bright-field images are shown in (**b**), with corresponding fluorescence images in (**c**). The latter also revealed a saturated GRP region between the droplet boundary and the contact line. (**d**) Evaporation of 32 μM tick-GRP77 on a surface-passivated glass slide showing numerous spherical tick-GRP77 condensates. (**e**) Two similar-sized condensates coalescing and relaxing into a bigger condensate over a time span of a few seconds. (**f**) A typical example showing the change in the aspect ratio as a function of time during a fusion event. The blue line shows an exponential decay fit (R^2^ = 0.99), giving the fusion timescale, τ ≈ 3 s. (**g**) A linear fit for the decay time (τ) against the characteristic length scale (*l*) for several fusion events (n = 14) gives the inverse capillary velocity (η/γ) of 0.92 s/μm (R^2^ = 0.47). (**h**) The phase-separated solution eventually dries out forming salt crystals. However, rehydration of the dried sample with pure water completely resolubilizes the salt crystals as well as the condensates. Evaporation of the resolubilized sample again leads to LLPS demonstrating the reversibility of the phase separation process. For fluorescence imaging, the samples were doped with 5 mol% OG488-GRP77.

A rapid non-homogenous distribution of tick-GRP77 due to the coffee-ring effect was evident in fluorescence imaging, along with intense fluorescence intensity in the region between the contact line and the droplet boundary, indicating a high local concentration of the protein (**Figure 2c**). We defined t_*i*_ as the time interval between droplet deposition and contact line pinning. Interestingly, within a minute after the contact line was established, we observed the sudden appearance of numerous micron-sized droplets near the contact line (**Figure 2b-right**), hinting at LLPS of tick-GRP77 by surpassing the critical concentration. The GRP-rich nature of these droplets was evident from the corresponding fluorescence images (**Figure 2c-right**). Repeating the evaporation assay on a surface-passivated glass slide with 5% w/v polyvinyl alcohol (PVA; see Materials and Methods for details) gave a clearer picture of micron-sized, spherical tick-GRP77 condensates (**Figure 2d**).

The region between the contact line and droplet boundary is an inverted phase, where the continuous phase is the protein-rich condensed phase, interspersed with dilute phase droplets. This likely happens because this very thin region rapidly accumulates a high-volume fraction of the protein, and has been observed for other evaporating condensate samples^45^. Further confirmation of this dense continuous phase comes from the fact that we observed fusion of tick-GRP77 condensates with the inverted phase, and a closer look also revealed GRP-depleted dilute phase droplets (**Supplementary Figure 3**), similar to earlier reports^45^.

As coacervation is a concentration-dependent process, one would expect the time until the onset of coacervation to be dependent on the initial protein concentration. Indeed, increasing the initial protein concentration from 16 to 64 μM gradually decreased the time required for onset of coacervation from 20 min to 14 min, at constant initial salt concentration (PBS, pH 7.4; **Supplementary Figure 4**). The onset time for higher (128 μM) initial concentration did not decrease further. To further validate the tick-GRP77-specific nature of the formed condensates, we performed several negative controls (**Supplementary Figure 5**): Evaporation of tick-GRP77 solubilized in 140 mM NaCl led to similar condensate formation. However, tick-GRP77 solubilized in pure water did not lead to the formation of condensates. This suggested a clear role of salts in promoting phase separation of tick-GRP77, as will become clear in later sections. Using bovine serum albumin (a globular protein without any disordered regions) in PBS buffer did not form any condensates but rather formed only salt crystals. Similarly, evaporation of only buffer solution (PBS at pH 7.4) did not result in any kind of LLPS behavior.

One of the hallmark properties of condensates is their ability to form spherical droplets which eventually coalesce and relax into a bigger droplet. This is due to the surface energy minimization and the rapid reorganization of molecular interactions within the liquid droplet^23, 46^. We readily observed numerous fusion events between tick-GRP77 condensates upon physical contact with each other, followed by their relaxation into a bigger spherical condensate (**Figure 2e; Supplementary Movie 2**). Tracking the aspect ratio (major to minor axis ratio) of the fusing droplets showed an exponential relaxation to a sphere^47, 48^. **Figure 2f** shows a typical example of the evolution of aspect ratio with a fitted exponential decay (*A* = 1 + 1.15e^−0.36t^; *R*^2^ = 0.99), giving a relaxation coefficient τ ≈ 3 *s* (*n* = 14). Following the relation 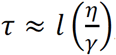, where *l* is the average radius of the fusing droplets, η is the viscosity of the droplet, and γ is the surface tension, a plot of τ against *l* gave us an inverse capillary velocity 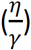 of 0.92 s/μm(**Figure 2g**; *n* = 14; *R*^2^ = 0.47; see Materials and Methods for details), implying micrometer-sized tick-GRP77 condensates behaved like liquid on a timescale longer than a second^48^.

Another key property of LLPS is its reversible nature, at least over short time scales. We checked this by repeating the above-mentioned droplet evaporation assay (32 μM GRP in PBS, pH 7.4) but then immediately rehydrating the sample to see its effect on the condensates. As expected, we observed the formation of GRP condensates and after complete evaporation, the sample crystalized due to the presence of salts (**Figure 2h**). However, upon rehydrating the crystalized sample with 2 μL of pure water, we immediately observed complete re-solubilization not only of the crystals but also of the GRP condensates. This clearly demonstrated the reversible nature of the LLPS process. Upon the second evaporation cycle, GRP condensates reformed in a similar manner. In conclusion, tick-GRP77 was observed to undergo simple coacervation and form viscous liquid droplets.

### The C-terminus of tick-GRP77 promotes phase separation via cation-pi and pi-pi interactions while the N-terminus prolongs LLPS through charge repulsion

To identify the protein regions responsible for phase separation and the underlying intermolecular interactions, we synthesized two distinct fractions of tick-GRP77: a 32-amino acid-long N-terminus (20–51) and the remaining 45-amino acid-long C-terminus (52–96), as depicted in **Figure 3a**. Both fractions are predicted to be disordered (**Figure 1d**) and have comparable glycine contents, 9 residues (28%) and 11 residues (24%) respectively. One clear difference is that the N-terminus is rich in acidic amino acids, giving it a net negative charge of -3.4 at pH 7.4. On the contrary, the C-terminus is relatively rich in basic amino acid residues giving it a net positive charge of 2.5 at pH 7.4 (**Figure 3b**). Conducting droplet evaporation assays for both fractions under identical conditions (50 μM solutions in PBS, pH 7.4) revealed a significant difference between the onset time of coacervation for the two termini. The N-terminus showed a similar timescale as tick-GRP77 with respect to the onset of coacervation, with a *t*_*i*_ ≈13 minutes (condensates at *t* = 14 min are shown in **Figure 3c**). On the other hand, the C-terminus underwent phase separation much earlier, with a *t*_*i*_ ≈ 3 minutes (condensates at *t* = 4 min are shown in **Figure 3d**). C-terminus condensates showed a tendency to wet the untreated glass surface (**Figure 3d**) but passivating the surface with 5% w/v PVA clarified the formation of spherical condensates (**Figure 3e**). Also, OG488-GRP77 (5 mol%) readily partitioned within the condensates, revealing their protein-rich interior.

**Figure 3:**
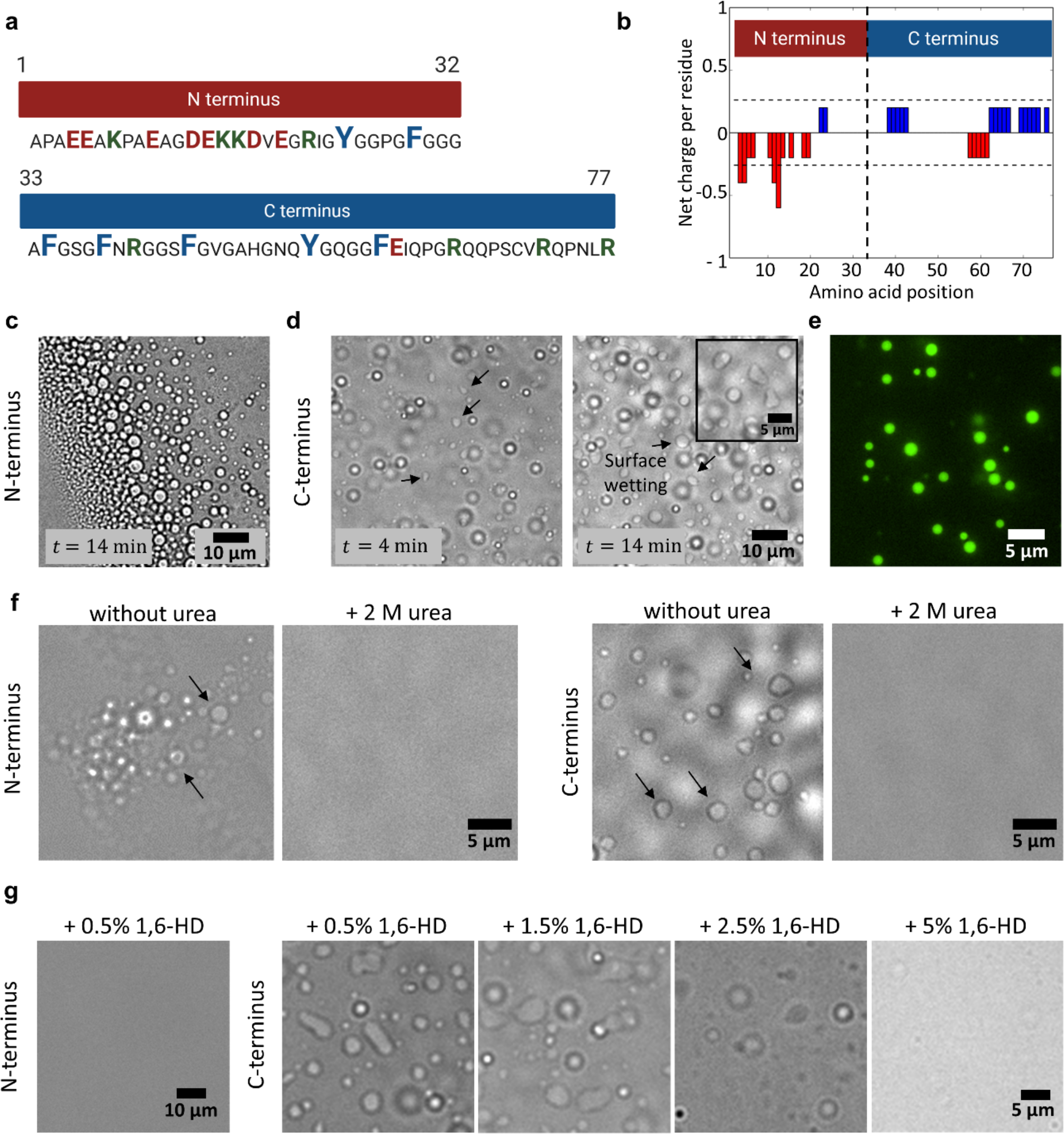
The C-terminus of tick-GRP77 is the main driver of phase separation. **(a)** Two distinct fractions of GRP77, the N-terminus (19–50) and C-terminus (51–96) have a similar glycine content but varying basic (green), acidic (red) and aromatic (blue) amino acid residues. (**b**) Net charge per residue as a function of amino acid position for the two termini. The N-terminus is strongly negatively charged while the C-terminus is moderately positively charged. (**c**) Droplet evaporation assay of the N-terminus leads to phase separation and formation of micron-sized condensates on a similar time scale as that of GRP77 (*t*_*i*_ ≈ 13 min). (**d**) Droplet evaporation assay of the C-terminus leads to much quicker phase separation (*t*_*i*_ ≈ 3 min). The formed condensates readily wet the glass surface. (**e**) Performing an evaporation assay on PVA-passivated glass slide prevented surface wetting leading to the formation of spherical C-terminus condensates. OG488-GRP77 (5 mol%) readily partitioned in the condensates. (**f)** Both N- and C-terminus condensates formed via droplet evaporation assay dissolved upon addition of 0.3 μL of 2 M urea indicating active role of hydrogen bonding in LLPS formation. (**g**) N-terminus condensates formed via evaporation dissolved upon addition of 0.5% w/v 1,6-HD (left). On the contrary, C-terminus condensates remained unaffected in the presence of 1.5% w/v 1,6-HD, and completely dissolved only at 5% w/v, indicating a prominent role of hydrophobic interactions compared to N-terminus condensates. Starting concentrations for both termini were 50 μM in PBS, pH 7.4 in all the experiments.

Apart from electrostatic interactions, π-π, cation-π, hydrophobic, and hydrogen bonding also play an important role in the formation of condensates^21^. **Figure 3a** shows that the N-terminus contains four cationic (three lysine and one arginine) and two aromatic (one tyrosine and one phenylalanine) residues. On the other hand, the C-terminus contains four cationic (all arginine) and five aromatic (four phenylalanine and one tyrosine) residues. Thus, the C-terminus can form extensive cation-π interactions compared to the N-terminus. Furthermore, the aromatic amino acids in the C-terminus are interspersed and separated by small groups of non-aromatic units which can be compared to the spacer-and-sticker model and capable of forming π-π interactions^49^. In addition to that, the five aromatic residues are fairly well-spaced and such equally spaced aromatic residues have been proposed to form liquid condensates^50^. Also, arginine (prominent in the C-terminus) has been proven to be relatively more hydrophobic than lysine (prominent in the N-terminus) due to the presence of π-electron-rich guanidium group which allows them to form π-π bonds along with cation-π bonds^51^. Finally, simple coacervation is more favorable if the protein contains relatively hydrophobic stickers and polar spacers^52^, which indeed seems to be the case for the C-terminus with multiple serine, asparagine, and glutamine residues. Earlier reports also suggest that arginine-glycine domains form cation-π interactions with phenylalanine and thus promote LLPS^53, 54^, which is the case at the C-terminus. Hence, it can be concluded that the C-terminus can readily form cation-π as well as hydrophobic interactions like π-π stacking explaining the rapid onset of coacervation. The observed delay in the initiation of coacervation for the N-terminus can be attributed to the lack of these interactions and the electrostatic repulsion between negatively charged amino acid residues. Thus, the C-terminus is the main promoter of tick-GRP77 LLPS while the high solubility and charge repulsion at the N-terminus delays the LLPS process and can act as a regulator.

We further investigated the role of inter- and intramolecular forces in the coacervation process. We first tested the role of hydrogen bonding by using urea, which efficiently forms hydrogen bonds with the amide moieties^55^. We used the droplet evaporation assay for both termini (50 μM in PBS at pH 7.4) and added 2 M urea solution (0.3 μL to ∼1 μL droplet, so final urea concentration of ∼0.5 M) as soon as the condensates were formed. Microscopic visualization showed both N- and C-terminus condensates immediately dissolved upon urea addition (**Figure 3f**) indicating an active participation of hydrogen bonding between the peptide backbone of GRP as well as amino acid residues such as histidine and tyrosine^56^. To check the role of hydrophobic forces in driving LLPS, we exposed N- and C-terminus condensates (50 μM solutions in PBS at pH 7.4) to 1,6-hexanediol (1,6-HD) which is widely used to dissolve liquid condensates, likely by disrupting the hydrophobic interactions involved in protein-protein interactions^57^. We again utilized the droplet evaporation assay and 1,6-HD was added as soon as the condensates formed. We observed that N-terminus condensates dissolved already after addition of 0.5% w/v 1,6-HD (0.3 μL 1,6-HD added to a ∼1 μL droplet, so final 1,6-HD concentration was ∼10 mM; **Figure 3g: left**). On the contrary, C-terminus condensates remained unaffected until 1.5% w/v 1,6-HD (final 1,6-HD concentration ∼40 mM) and dissolved completely only at 5% w/v (final 1,6-HD concentration ∼130 mM) (**Figure 3g: right**). These results confirm our previous conclusion that the C-terminus phase separates via extensive hydrophobic interactions and thus requires more 1,6-HD to disrupt sufficient condensate-forming interactions.

Based on the independent investigations of N- and C-termini, we conclude that cation-π and π-π stacking of aromatic amino acids^62^ are important for the phase transitioning of tick-GRP77.

### Tick-GRP77 forms liquid condensates in presence of kosmotropic salts

Tick saliva is salty and hygroscopic which helps ticks to absorb moisture to stay hydrated^58^. Additionally, tick saliva also contains enzymes like apyrase^59^ and acid phosphatase, which can increase local phosphate concentration through adenosine triphosphate (ATP) degradation. Studies have reported that kosmotropic ions (HPO_4_^2-^, SO_4_^2-^, etc.) facilitate LLPS^60, 61^ by having strong bonding interactions with water molecules, thereby decreasing protein solubility, and promoting phase separation. Given our previous observation that pure GRP solution did not undergo LLPS and to test the effect of kosmotropic salts on tick-GRP77 phase transition, we chose disodium hydrogen phosphate (Na_2_HPO_4_) salt, mimicking the potential inorganic phosphate release in saliva. Indeed, addition of 1 M Na_2_HPO_4_ to 63 μM and 125 μM tick-GRP77 solutions led to instant coacervation. We tested several different GRP concentrations and incubation times up to several hours to obtain a phase diagram (**Figure 4a**). As can be seen, a short incubation time of 1.5 hours at room temperature (22 °C) was enough to form condensates also at much lower concentrations (16 μM and 31 μM). Concentrations below 16 μM did not result in condensation even after 5.5 hours. At lower Na_2_HPO_4_ concentrations (0.5 M), we did not observe immediate condensation, but only after several hours of incubation and at high GRP concentrations (>63 μM), did condensates emerge (**Supplementary Figure 6).**

**Figure 4:**
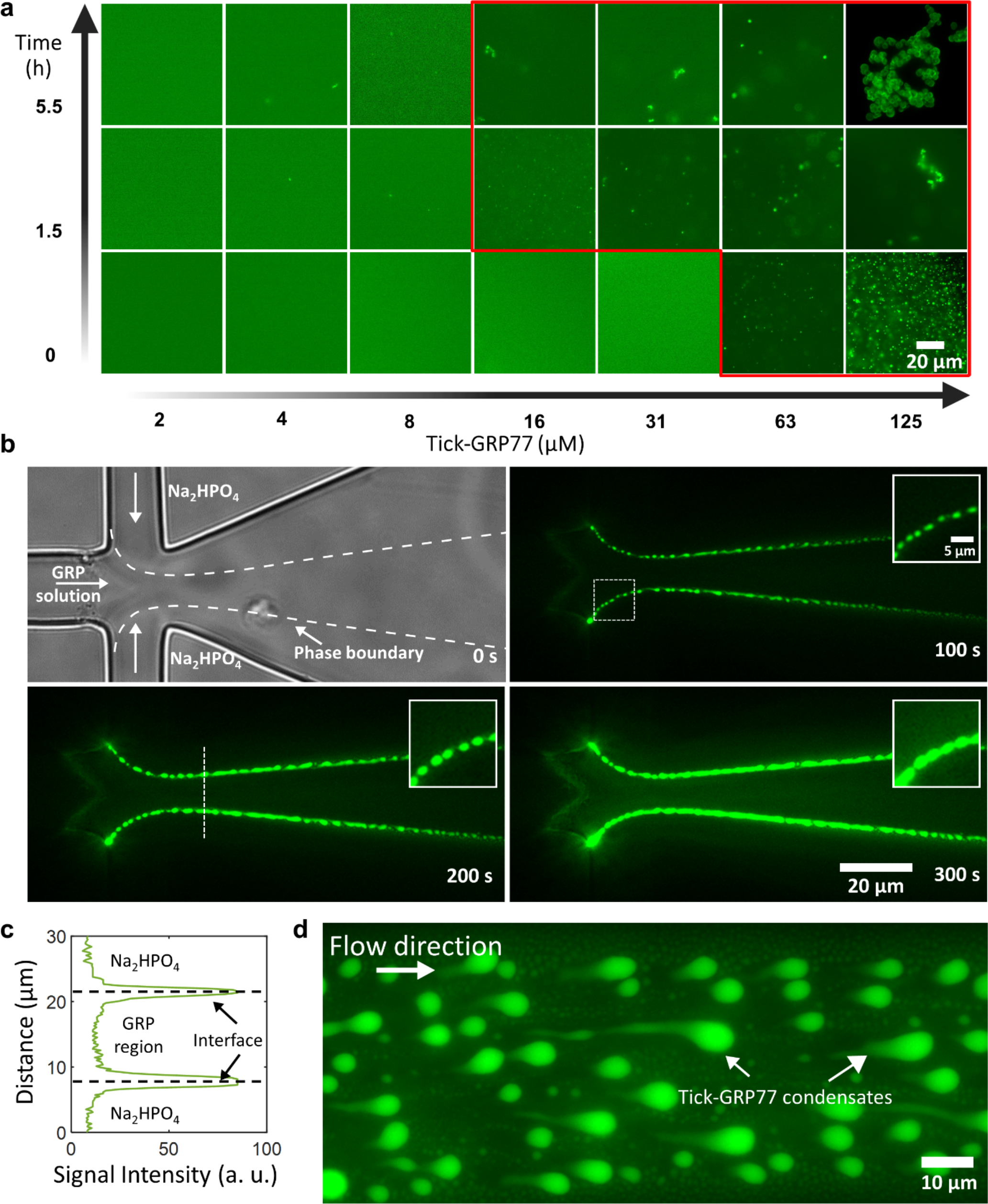
Tick-GRP77 forms condensates in presence of kosmotropic salts. **(a)** A phase diagram showing LLPS behavior of tick-GRP77 in presence of 1 M Na_2_HPO_4_ as a function of protein concentration and incubation time. The red region indicates the condensation regime; concentrations above 63 μM resulted in instant coacervation, whereas for lower GRP concentrations (16 and 31 μM), phase separation was observed after 1.5 h incubation at room temperature. (**b**) Left top: Bright-field image showing a flow-focusing junction of the microfluidic device. The inner stream containing 63 μM tick-GRP77 in pure water is focused by the two streams of 2 M Na_2_HPO_4_. As the three streams flow side-by-side, GRP condensates are observed to form exclusively at the GRP-salt interface. The condensates wet the channel wall and become bigger over time. (**c**) A fluorescence intensity plot (corresponding to the dotted line in b) show the intensity profile across the interface of GRP-salt streams. (**d**) GRP condensates stuck to the channel walls downstream of the junction are deformed into tear-shaped droplets because of the fluid flow, indicating their liquid-like nature.

To further clarify the instant nature of the coacervation and the role of kosmotropic salts, we performed flow-focusing experiments using microfluidic devices. A key feature of microfluidic systems is their laminar flow^62^ owing to the low Reynolds number^63^, allowing strictly diffusion-based mixing between the co-flowing fluid streams. We used a lab-on-a-chip flow-focusing device, allowing the GRP and salt streams to meet and co-flow together without mixing (see Materials and Methods for details; **Supplementary Figure 7**). GRP solution (63 μM in pure water; 5 mol% OG488-GRP77) was injected in the inner aqueous channel and 2 M Na_2_HPO_4_ solution was injected in outer aqueous channels, forming two sharp tick-GRP77-Na_2_HPO_4_ interfaces at the junction (**Figure 4b; left top**). We observed immediate formation of tick-GRP77 condensate droplets at the interface, which adhered to the channel walls and continuously increased in their size as more and more condensate phase got accumulated (**Figure 4b; fluorescence panels; Supplementary Movie 3**). A line profile perpendicular to the interface shows two clear fluorescence intensity peaks at the interfaces, clarifying the salt-induced condensation (**Figure 4c**). The liquid nature of these condensates became more evident further down the channel (approximately 500 μm downstream of the junction) where the condensate droplets wetting the channel walls were deformed by the fluid flow into tear-shaped droplets (**Figure 4d**). This made it clear that the formed structures are not protein aggregates or precipitates, but phase-separated liquid droplets.

### Tick-GRP77 condensates undergo liquid-to-gel transition

Natural tick saliva eventually undergoes a liquid-to-solid phase transition to form a hard cement cone. We observed several instances in our experiments which indicate that LLPS may be an intermediate stage in this transition. Conducting the droplet evaporation assay using high initial tick-GRP77 concentration (initial concentration 500 μM in PBS, pH 7.4) on a hydrophobic glass slide not only led to condensate formation, but the formed condensates fused together to form network-like structures (**Figure 5a; Supplementary Movie 4**). A time-lapse showing an example of a fiber formation leading to an interconnected network of tick-GRP77 condensates can be seen in **Figure 5b**. On similar lines, a network composed of stretched sheets and fibers adhering to a hydrophilic glass surface (coated with PVA) was obtained during evaporation experiments, as shown in **Figure 5c**. These examples point to the transition of liquid condensates into viscoelastic gel-like networks on solid surfaces, irrespective of their hydrophilic/phobic nature.

**Figure 5:**
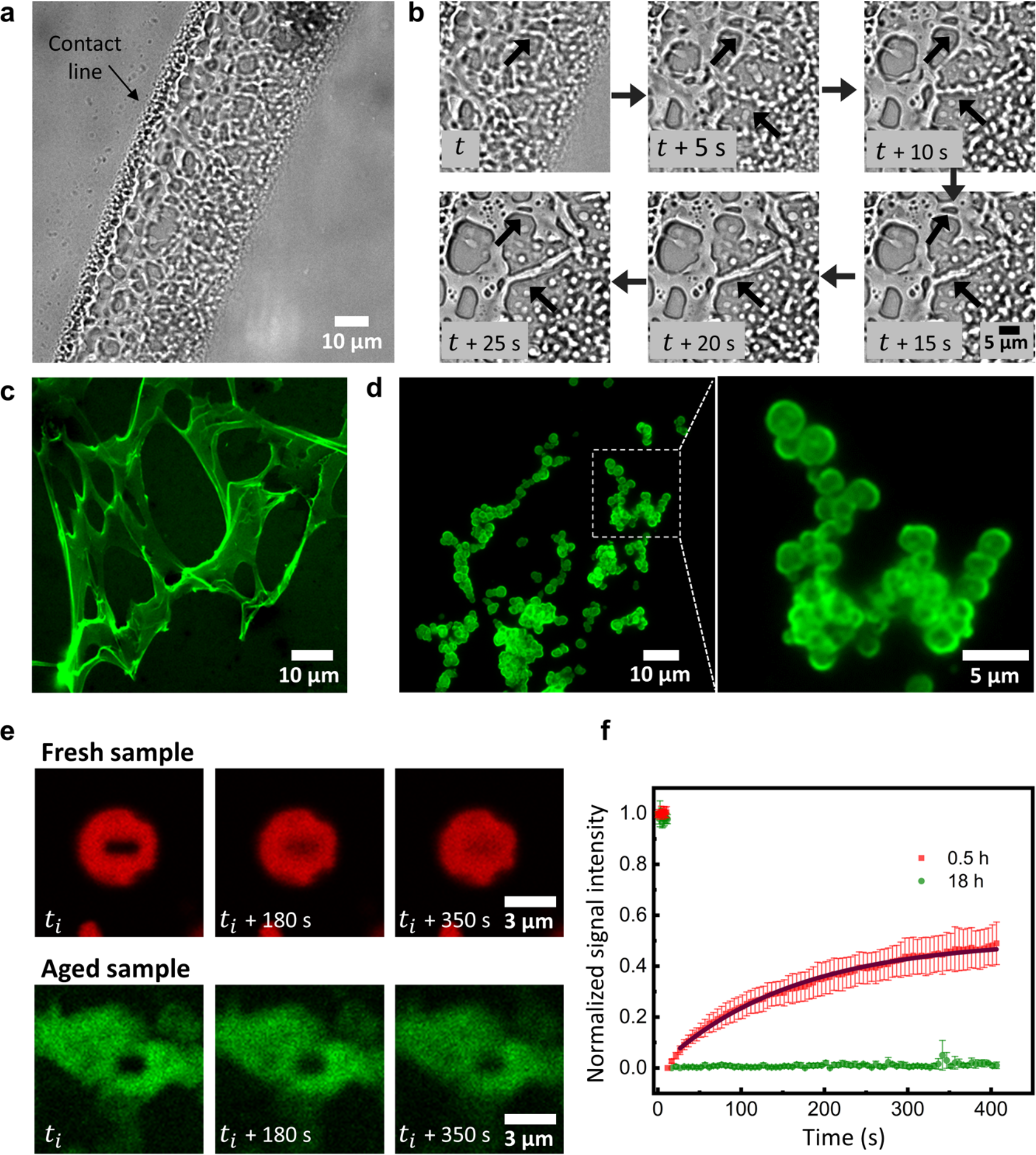
GRP condensates form viscoelastic networks and solid-like aggregates at higher concentrations and longer incubation times. **(a)** Droplet evaporation assay of a concentrated tick-GRP77 solution (500 μM) resulted in the formation of condensate fibers, leading to interconnected gel-like networks. **(b)** Time-lapse showing the assembly of the condensates into fiber-like structures. Two such events are indicated by black arrows. **(c)** Fluorescence image of an interconnected stretched network formed during evaporation assay of tick-GRP77 (32 μM) solubilized in 5-times concentrated PBS (pH 7.4) on a PVA-passivated hydrophilic glass slide. (**d**) Incubating tick-GRP77 solution (125 μM) in 1 M Na_2_HPO_4_ for 5.5 hours at room temperature formed stable condensate clusters, with complete arrest of coalescence. The zoom-in shows individual condensates physically connected to each other without undergoing fusion. (**e**) Time-lapse showing the fluorescence recovery of GRP condensates (125 μM) for fresh (0.5 h) and older (18 h) samples. The condensates were formed in presence of 1 M Na_2_HPO_4_ salt. (**f**) Fluorescence recovery curve showing that the freshly formed condensates have a higher fraction of the mobile phase compared to matured GRP condensates. The black line shows exponential fit to the dataset (R^2^ = 0.99). Samples in c, d, and e were doped with 5 mol% OG488-GRP.

More solid-like structures were observed during salt-induced coacervation when using a high concentration of GRP77 (125 μM) over longer incubation times (5.5 hours). Here, the coalescence of the condensates was clearly arrested, resulting in stable grape-like clusters (**Figure 5d**). To verify the solidification or the ageing of GRP condensates over time, we performed fluorescence recovery after photobleaching (FRAP) experiments (tick-GRP77 concentration = 125 μM, Na_2_HPO_4_ salt concentration = 1 M). A comparative FRAP study was conducted on freshly prepared (0.5 hours after preparation) (**Figure 5e top; Supplementary Movie 5**) condensates and older samples (18 hours of incubation) as shown in (**Figure 5e bottom; Supplementary Movie 6).** We photobleached a small region within the condensates (see Material and Methods for details) and recorded the recovery of the fluorescent intensity (**Figure 5e**). We quantified the fluorescence recovery over time for both freshly prepared (*n* = 5) and aged (*n* = 4) samples which revealed two important findings (**Figure 5f**). First, the post-bleaching fluorescence intensity did not show full recovery in both the cases which shows the arrested motion of GRP molecules within condensates already in early stages and their apparent viscoelastic behavior. Secondly, fresh samples showed much more recovery (≈49%) compared to the aged samples (≈2%). This drastic decrease in the recovery of fluorescent intensity for the aged sample shows further reduction in the fraction of mobile phase over time and the transformation from liquid-to-solid-like state. We fitted the normalized intensity *I(t)* with an exponential function, 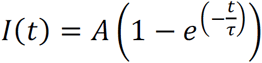, where *A* and τ indicate the amplitude of recovery and the relaxation time, respectively^48, 64^. For the fresh sample, we obtained a relaxation time, *τ_fresh_* = 160 s, with the diffusion coefficient of OG488-GRP (*D_app_*) in the order of ∼2.2 × 10^−3^ μm^2^/s (see Material and Methods for details). Using Stokes-Einstein relation 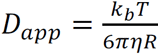, where *k_b_T* is the thermal energy scale, η is the droplet viscosity, and *R* is the hydrodynamic radius of OG488-GRP77 (∼2.5 nm for an unfolded 7.8 kDa protein^65^), we estimated the condensate viscosity to be ∼42 Pa.s which is similar to that of ketchup and mustard. Plugging the value into the inverse capillary velocity 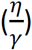 led to the estimation of the interfacial tension ∼46 μN/m, similar to the very low values reported for macromolecular liquids^66^ as well as protein condensates^47^. Thus, microscopic and FRAP analysis together demonstrated tick-GRP77 condensates to be highly viscous liquids with ultralow interfacial tension, capable of forming viscoelastic networks and exhibiting ageing over the course of a few hours. This liquid-to-gel transition is highly relevant to the tick cement cone formation that also takes place over several hours^10^.

### Natural tick saliva shows evidence of protein-rich biomolecular condensates

All the above-mentioned *in vitro* experiments indicated a strong phase separation tendency of tick-GRP77, which encouraged us to further investigate if natural tick saliva exhibits similar phase separating behavior. We collected ticks from the forest region of Veluwe, The Netherlands, in the months of June and July 2022, when the temperature is typically around 20 °C. In these conditions, ticks “quest” by climbing up grass and low-lying vegetation and waiting for a potential host to pass by, which they will grab and climb on to. We collected ticks from the grass and small shrubs by using a tick-dragging method as previously described^67^. We collected ticks belonging to the species *Ixodes ricinus* at different life stages: nymphs, adult males, and adult females (**Supplementary Figure 8**). The adult female salivary glands of the *Ixodidae* groups consists of type I, II and III acini cells^58, 68^, among which the type II acini cells are associated with cement formation^68, 69^. We dissected adult females, isolated the salivary glands^70^, and extracted the contents of salivary glands using sonication (**Figure 6a**; see Materials and Methods for details). An important factor to note here is that the ticks we collected were not blood-fed and thus likely have a lower expression of GRPs compared to blood-fed ticks^71^. Additionally, it has been reported that ticks species with long hypostome like *Ixodes ricinus* have lower level of cement production^72^, and thus likely lower expression of glycine-rich proteins^16^. Nonetheless, microscopic visualization of the salivary gland extract revealed numerous micron-sized spherical droplets (**Figure 6b**). Moreover, we also recorded a droplet fusion event indicating their liquid nature (**Figure 6c**). Importantly, when we doped the saliva sample with fluorescently labelled OG488-GRP, we observed strong partitioning of OG488-GRP in these droplets (**Figure 6d**). This clearly rules out the possibility of these droplets being lipid or fat droplets and at least hint at their protein-rich nature, and likely GRP-rich. In addition to the droplets, we also observed GRP-rich fiber-like structures after incubation (one hour) at room temperature, hinting to liquid-to-solid transition and corroborating with our *in vitro* findings (**Figure 6e**).

**Figure 6:**
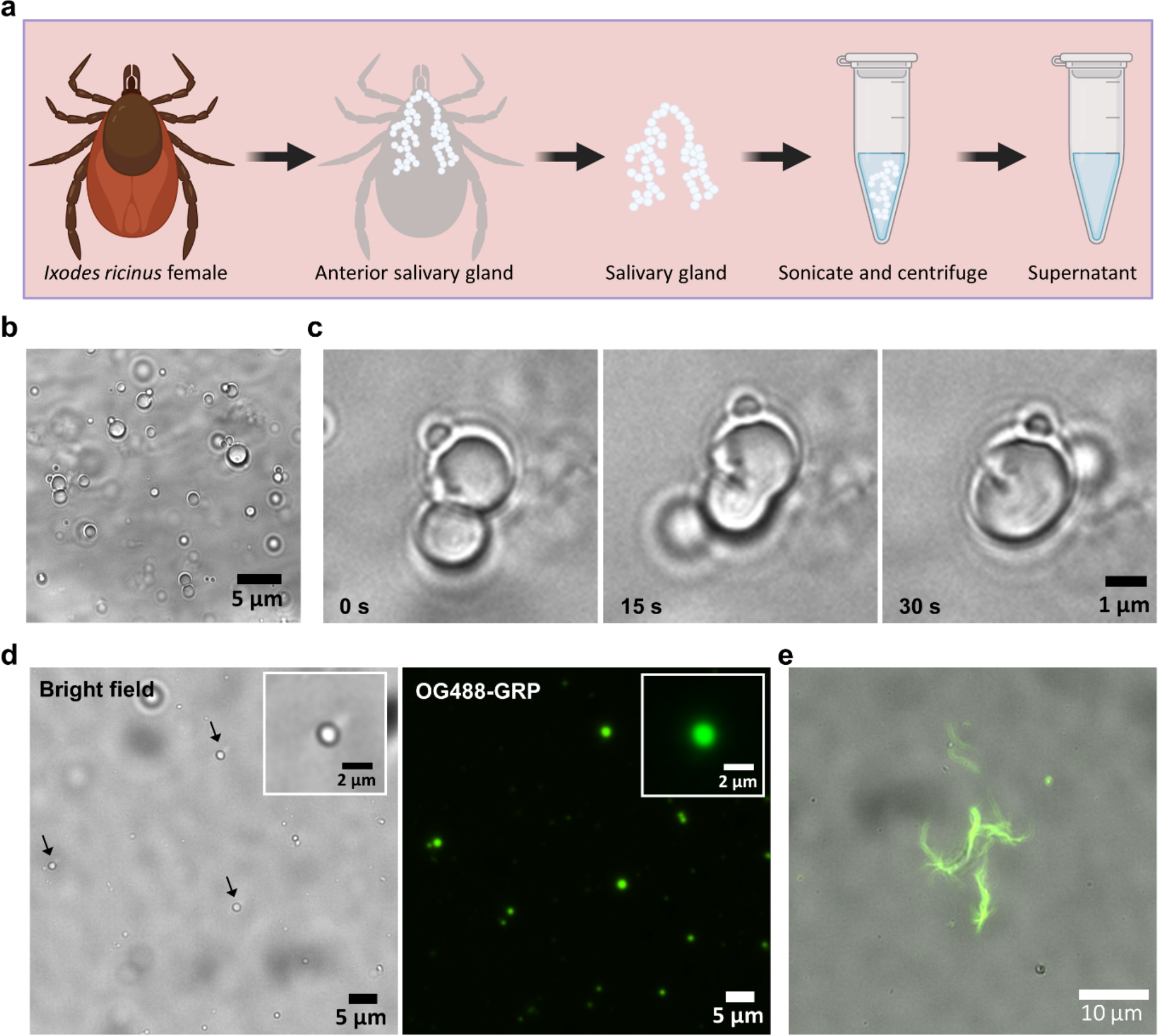
Saliva extracted from ticks hints at the presence of protein-rich phase-separated droplets. **(a)** Schematic showing the extraction of the salivary gland from a non-blood-fed tick (Ixodes ricinus). The collected supernatant was used in the following experiments. (**b**) Bright-field visualization of the supernatant showing numerous micron-sized spherical droplets. (**c**) Fusion of two droplets, indicating their liquid nature. (**d**) Spiking the supernatant with fluorescently labelled GRP77 (OG488-GRP77) readily led to its partitioning inside the phase-separated droplets indicating their protein-rich nature and possible GRP-rich interior. (**e**) Overlay of bright-field and fluorescence images showing fiber-like structures observed in the supernatant after approximately one hour, indicating liquid-to-solid transitioning of the droplets. OG488-GRP77 showed strong partitioning in these structures.

## Conclusion

We have demonstrated that the glycine-rich protein (GRP) present in tick saliva undergoes liquid-liquid phase separation and exhibits ageing behavior to form viscoelastic gel-like structures. We showed that GRP condensates form via simple coacervation, and their material properties change over a time frame of a few hours. Considering that tick cement deposition and solidification is a slow process^12^, our results suggest a plausible role of GRPs in the cement cone formation, namely to induce liquid-liquid and subsequent liquid-to-solid phase transition, a vital process in the attachment of ticks to their host.

The phase separation of tick GRP is induced via multiple weak interactions. The negatively charged N-terminus keeps the protein soluble while a chain of regularly spaced aromatic residues separated by glycine-rich residues promote cation-π and π–π interactions leading to coacervation. Additionally, hydrogen bonding and hydrophobic interactions, especially given the glycine-rich backbone, also play a significant role^73^.

GRPs are also found in adhesive proteins from various other organisms such as mussels, sandcastle worms, and velvet worm^2, 31, 32, 34, 73^. Thus, understanding the phase transitions of GRPs may reveal possible common unifying principles in bioadhesive proteins functioning in distinct environments across diverse animal species^73^. In future, it will be interesting to explore the role of other relevant parameters, like pH and temperature, on the phase transition behavior of tick GRPs. Given the fact ticks detach from the host post blood-feeding, a dissolution mechanism of the hardened cement cone will also be worth investigating^17, 74^.

Tick GRPs could potentially be used for the development of medical sealants like tissue glues because of their biodegradable and biocompatible properties^75^. The role of GRPs in host adhesion may provide insights to manage ticks and tick-borne pathogens, a major problem worldwide, and particularly in developing regions in tropical regions, especially due to the lack of sustainable control methods^76^. Developing chemicals that can interfere with the phase transition of tick-GRP77 and thus inhibit cement cone formation may provide an effective solution. Anti-tick vaccines are a promising tick control strategy with the potential to provide long-term protection^77, 78, 79^. Characterization of GRPs have shown they are immunogenic, play a role in host’s immune system evasion, and are required for successful tick attachment, making them promising anti-tick candidates^80, 81, 82^. Also, condensates have a high potential to be effective drug delivery systems owing to their rapid and aqueous self-assembly, possibility of targeting, and enhanced bioactivity of the cargo^83^. Thus, testing the potential of GRPs as novel anti-tick vaccine agents could be another outcome of this research.

## Materials and Methods

### GRP Synthesis and fluorescent labelling

#### Boc-based solid-phase peptide synthesis (SPPS)

The N- (H_2_N-APAEEAKPAE AGDEKKDVEG RIGYGGPGFG GG-MPAL) and C- (H_2_N-AFGSGFNRGG SFGVGAHGNQ YGQGGFEIQP GRQQPSCVRQ HPNLR-OH) terminal peptide fragments of the protein were synthesized on 0.10 mmol scale. 0.18 g of resin was used for both syntheses, with 4-(Hydroxymethyl) phenylacetamidomethyl (PAM) polystyrene resin with preloaded leucine (Leu) and arginine (Arg) or the N- and C-terminal peptide fragments, respectively. The first alanine residue (Ala) of the C-terminal peptide fragment sequence was replaced by a cysteine residue (Cys) (A33C mutation) for NCL. Each amino acid was activated with 0.5 M 2-(6-Chloro-1-H-benzotriazole-1-yl)-1,1,3,3-tetramethylaminium hexafluorophosphate (HCTU) in *N,N*-dimethylformamide (DMF) and *N, N*-Diisopropylethylamine (DIPEA) before coupling. The coupling time for all amino acids was 10 minutes, except for serine (Ser), threonine (Thr), Arg, and asparagine (Asn) for which the coupling time was 20 minutes. For glycine (Gly), the coupling time was also set to 20 minutes for the N-terminal peptide fragment, whereas 10 minutes double couplings done for the C-terminal peptide fragment. After each coupling, the resin was washed with DMF, then treated with Trifluoroacetic acid (TFA) two times for 1 minute, and washed again with DMF. After glutamine (Gln) coupling (C-terminal peptide fragment synthesis), the resin was washed with DCM as well before and after TFA treatment to prevent intramolecular pyrrolidone formation. For thioester synthesis at the N-terminal peptide fragment, 3-mercaptopropionic acid (MPA) was coupled via a Leu residue and subsequent trityl group-deprotection was performed with a 95%/2.5%/2.5% TFA/Triisopropylsilane (TIS)/H_2_O mixture. The N- and C-terminal peptide fragments (495 mg and 440 mg, respectively) were deprotected and cleaved from the solid-phase by anhydrous hydrogen fluoride (HF) treatment for 1h at 0°C using 4% v/v *p*-cresol as scavenger. The peptides were precipitated in ice-cold diethyl ether, dissolved in a MeCN/H_2_O mixture containing 0.1% TFA, and lyophilized.

#### Native chemical ligation (NCL)

NCL of the unprotected synthetic peptide segments was performed as follows: 0.1 M TRIS buffer, pH 8, containing 6 M Gnd-HCl was added to dry peptides yielding approximately 10 mg/mL of peptide fragments. Subsequently, 1% v/v benzylmercaptan and thiophenol were added. The ligation reaction was performed in a heating block at 37 °C and the mixture was vortexed periodically to equilibrate the thiol additives. Reaction progress was analyzed with UPLC-MS. After the reaction was complete, thiophenol was removed by diethylether extraction of the reaction mixture (3 ×).

#### Desulfurization

Desulfurization was directly performed after the native chemical ligation reaction in order to convert the first Cys residue of the *C*-terminal peptide fragment into an Ala residue and thus to obtain the original GRP sequence. Desulfurization buffer was prepared by dissolving tris(2-carboxyethyl)phosphine (TCEP; 250 mM) in 5 mL 0.1 M TRIS buffer, pH 8, containing 6 M Gnd-HCl. The pH of the desulfurization buffer was adjusted by adding solid NaOH until it reached pH 7. Then, reduced L-glutathione (GSH; 40 mM) was added to 1 mL of this desulfurization buffer. 250 μL of this solution were added to the reaction mixture and VA-044 (6.25 mM) was added. The reaction was performed at 37°C and reaction progress was monitored using UPLC-MS until the product was observed. Analytical HPLC was performed to purify GRP by using a C18 column (150 mm × 4.6 mm) connected to a Prostar HPLC (Varian).

#### Acm deprotection

To remove the acetamidomethyl (Acm) protecting group of the cysteine residue in the *C*-terminal peptide fragment, the peptide was dissolved at a 2 mM concentration in 0.1 M TRIS, pH 7.25, containing 6 M Gnd-HCl. Then, 10 eq. Pd-Cl^2^ were added and the deprotection progress at 37 °C was monitored using UPLC-MS. After reaction completion, the formed Pd-complex was reduced with 20 mM DTT for 1 hour at 37 °C. Subsequently, the reaction mixture was purified on a C4 column (Vydac, 150 mm × 4.6 mm) with an appropriate gradient on the analytical HPLC system as described above. Measured monoisotopic mass: 7854.08 Da; calculated monoisotopic mass: 7853.735 Da.

#### In vitro experimentation and microscopic visualization

Stock solution of GRP was made in MilliQ water. Unless specified, all the evaporation experiments were performed in phosphate buffer saline (10 mM Phosphate buffer, 2.7 mM KCl, and 137 mM NaCl at pH 7.4). A 2 μl droplet of GRP solution was transferred on a coverslip (24 mm × 40 mm, Corning™ #1.5) and mounted on Nikon-Ti2-Eclipse inverted fluorescence microscope equipped with pE-300^ultra^ illumination system. For all the experiments, droplets were visualized using either Nikon Plan Apo 100× (numerical aperture, NA 1.45) oil objective or Nikon Plan Fluor 40× (NA 1.30) oil objective. For fluorescent visualization, GRP sample was doped with 5 mol% of OG488-GRP and excited via 482/35 nm excitation filter, 505 nm dichroic mirror and the emitted light was collected through 536/40 nm emission filter (Semrock). The samples were typically excited using 2-5% laser intensity and time-lapse images were acquired at exposure of 5-20 ms using a Prime BSI Express sCMOS camera. Confocal microscopy for visualizing the inverted phase was recorded using Nikon C2 CSLM, equipped with a 60× (NA 1.40) oil immersion objective. The GRP sample was doped with 5 mol% OG488-GRP and was imaged using 488 nm excitation laser, 560 nm LP dichroic mirror and 525/50 emission filter. We minimized the illumination intensity (1–2% of 15 mW) to prevent bleaching of the sample.

#### Surface functionalization of coverslips

Wherever required, coverslips were coated with 5% w/v solution of polyvinyl alcohol (molecular weight 30 -70 kDa, 87 – 90 % hydrolyzed) as described previously^85^. Briefly, the coverslip was plasma treated for 30 seconds at 12 MHz (RF mode high) using a plasma cleaner (Harrick plasma PDC-32G). A 10 μL drop of PVA solution was pipetted out in the center of the glass slide and allowed to rest for 5 min followed by gently removing of PVA by tilting the glass slide. To remove free PVA, the coverslip was gently washed with milli Q water. The glass slide was baked at 70 °C for 2 hours and stored at room temperature in a clean environment. To make the glass slide hydrophobic, ∼20 μL of (tridecafluoro- 1,1,2,2-tetrahydrooctyl)trichlorosilane was taken in a glass vial and placed in the desiccator along with glass slides. The set up was left under partial vacuum for ∼12 h to render the glass slides hydrophobic.

#### Droplet fusion experiments

A 2 μL sessile drop of tick-GRP solution (32 μM with 5% mol fraction of OG488-GRP; dissolved in PBS pH 7.4) was allowed to evaporate on a PVA-passivated glass slide. Droplet fusion events were recorded using 40× oil objective at time interval of 250 ms (exposure time of 10 ms) on Nikon Ti2 Eclipse fluorescence microscope. The aspect ratio of the fusing droplets was determined by fitting an ellipse using Fiji (Image J) and calculating aspect ratio 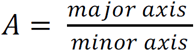 where major and minor axes are the long and short axes of the ellipse respectively. For analysis of fusing condensates, the change in aspect ratio with respect to time was plotted and the data was fit to the function of the form 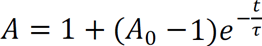; where t is the time, τ is the characteristic relaxation time, and *A*_0_ is the initial aspect ratio. Using the relation 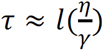, where *l* is the average diameter of the droplets, η is the viscosity and γ is the surface tension, the inverse capillary velocity 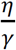 was calculated^48^.

#### Microfabrication

The master wafer was prepared according to the previously described UV lithography method^86^ and the protocol was adjusted to attain the channel height of 20 μm. To prepare the microfluidic device, polydimethylsiloxane (PDMS) and curing agent (SYLGARD™184 elastomer) were mixed in 10:1 weight ratio. The mixture was poured on the master, degassed using vacuum desiccator followed by baking at 70 °C for 4 hours. The hardened PDMS block was carefully removed, and inlets and outlet holes were punched using a biopsy punch of diameter 0.5 mm (Darwin microfluidics). The PDMS block was then bonded on a glass coverslip (Corning® #1) using a plasma cleaner (Harrick Plasma PDC-32G). The bonded device was baked at 80 °C for two hours and stored at room temperature. Elveflow pressure controller OB1-MK3 was used to flow GRP and salt solutions 2 M Na_2_HPO_4_ (10 mM Tris-Cl, pH 7). The fluid flow was maintained at constant pressure of 100 mbar and 20 mbar for the inner aqueous (GRP solution) and outer aqueous (Na_2_HPO_4_ solution) channels respectively.

#### Fluorescence recovery after Photobleaching

FRAP experiments were performed on Leica SP8-SMD microscope and 63× (NA 1.2) water objective. For bleaching, the region of interest (ROI) of approximately 1.5 μm length and 0.5 μm breadth was selected inside the condensates of approximately 5 μm in diameter. The ROI was bleached using 100% laser intensity for 2 seconds and recovery of the bleached area was recorded for every 5 seconds for approximately 6 minutes. Intensity of the bleached area was normalized using the equation, 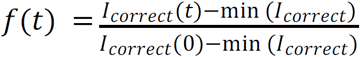, where 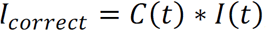, and 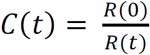. Here, *R*(*t*) and *I*(*t*) indicate the fluorescence intensity of the reference droplet at time *t* and the original fluorescence intensity of the bleached region at time *t*, respectively; min(*I_correct_*) indicates the minimum value of *I_correct_*, which is obtained right after the sample is bleached^64^. The normalized intensity was fitted using the function 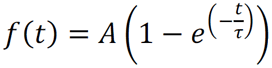, where *A* and τ indicate the amplitude of recovery and the relaxation time, respectively. The apparent diffusion coefficient (*D_app_*) was calculated using the formula 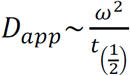 where 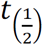 is the half-life fluorescence recovery and ω^2^ is the area of the bleached cross section. The half-life 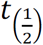 was calculated using the formula 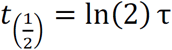.

#### Salivary gland extraction

Ticks of the species *Ixodes ricinus* were dissected according to a previously described protocol^70^, with the exception that the ticks were not blood-fed before salivary gland isolation. Salivary glands were resuspended in 100 μL milliQ water and their contents were extracted by mechanically disrupting the glands via sonication at 45 kHz for 5 minutes, followed by debris segregation via centrifugation at 13,000 g for 10 minutes. The supernatant was collected in a fresh tube and used for experimentation.

## Acknowledgements

We thank Jasper van der Gucht for his feedback on the manuscript. Elisa Nihoul is kindly acknowledged for her contribution to the GRP synthesis. We thank Hans Smid (www.bugsinspace.nl) for providing the images of ticks and Marcel Giesbers from Wageningen Electron Microscopy Centre for providing the scanning electron microscopy image of tick mouth parts. We acknowledge Niels Appelman for assisting with fabrication of microfluidic devices. S.D. acknowledges financial support from Dutch Research Council (grant number: OCENW.KLEIN.465). All the schematics were made using Biorender.com.

## Author contributions

K.G., P.T., and S.D. conceived the idea and designed the experiments. K.G., P.T., M.N., and C.C. performed *in vitro* experiments. K.G. and E.L.P. carried out tick collection and salivary gland extraction. K.G., P.T., M.N., C.C., and S.D. performed data analysis. D.S, S.B, and I.D. synthesized the proteins. K.G., P.T., M.N., I.D. and S.D. wrote the initial draft. K.G, M.N and S.D reviewed and edited the manuscript. S.D. acquired funding and performed project administration and supervision. All authors have read and agreed to the final version of the manuscript.

