## Supplementary Information for "Phase Separation and Ageing of Glycine-Rich Protein from Tick Adhesive"

**Supplementary Figures**

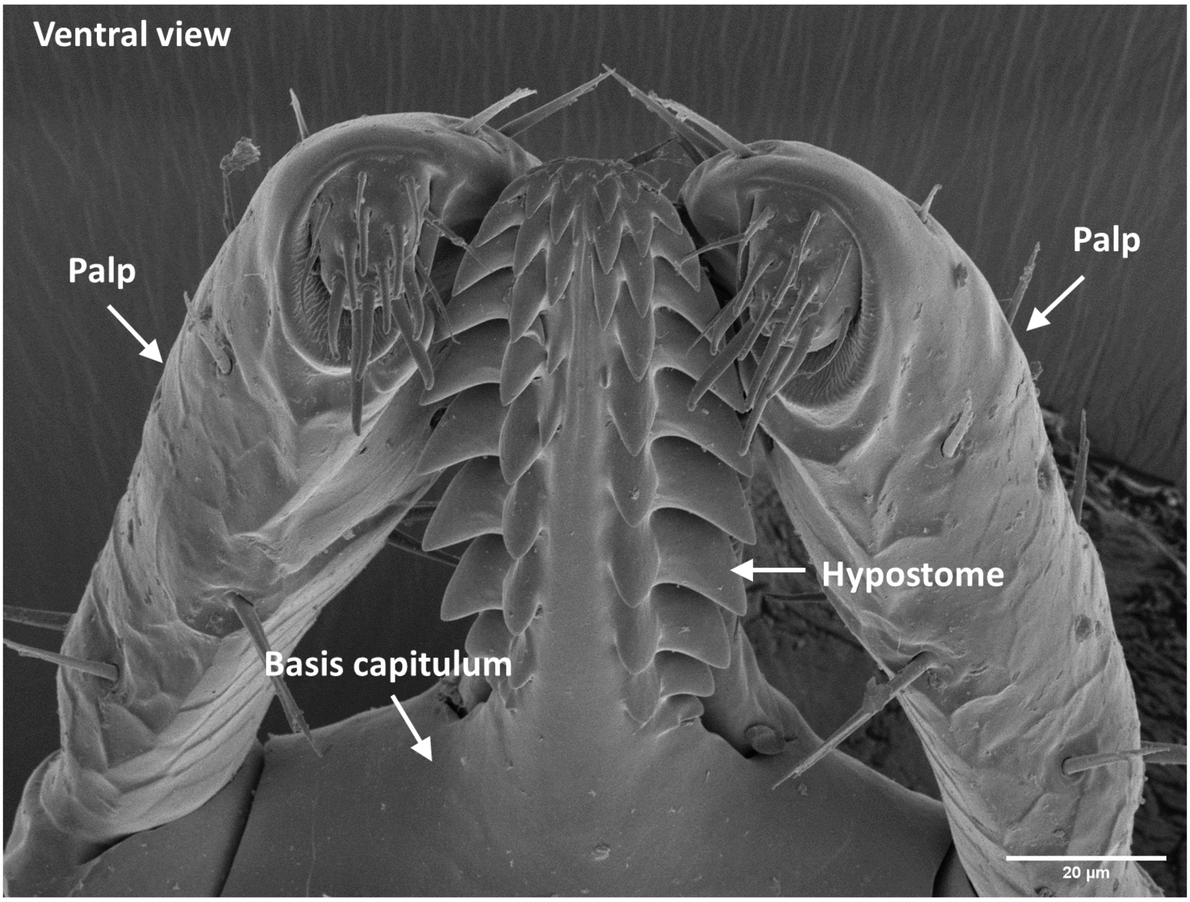


**Supplementary Figure 1. Tick mouth parts.** Electron microscopy of tick mouth parts visualized from the ventral side.

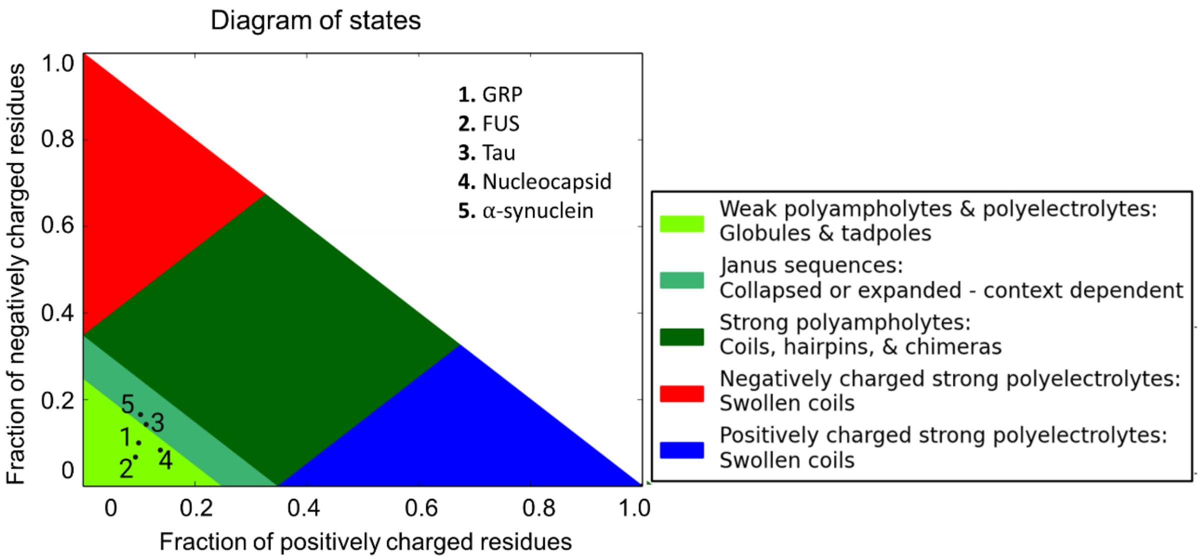


**Supplementary Figure 2. Tick-GRP77 falls in the same region within the diagram of states as many well-characterized condensate-forming proteins.** Diagram of states indicating tick-GRP77 as weak polyampholytes and polyelectrolytes and falls in close vicinity of FUS, tau, nucleocapsid protein and $\alpha$-synuclein, all of which are known to phase separate.

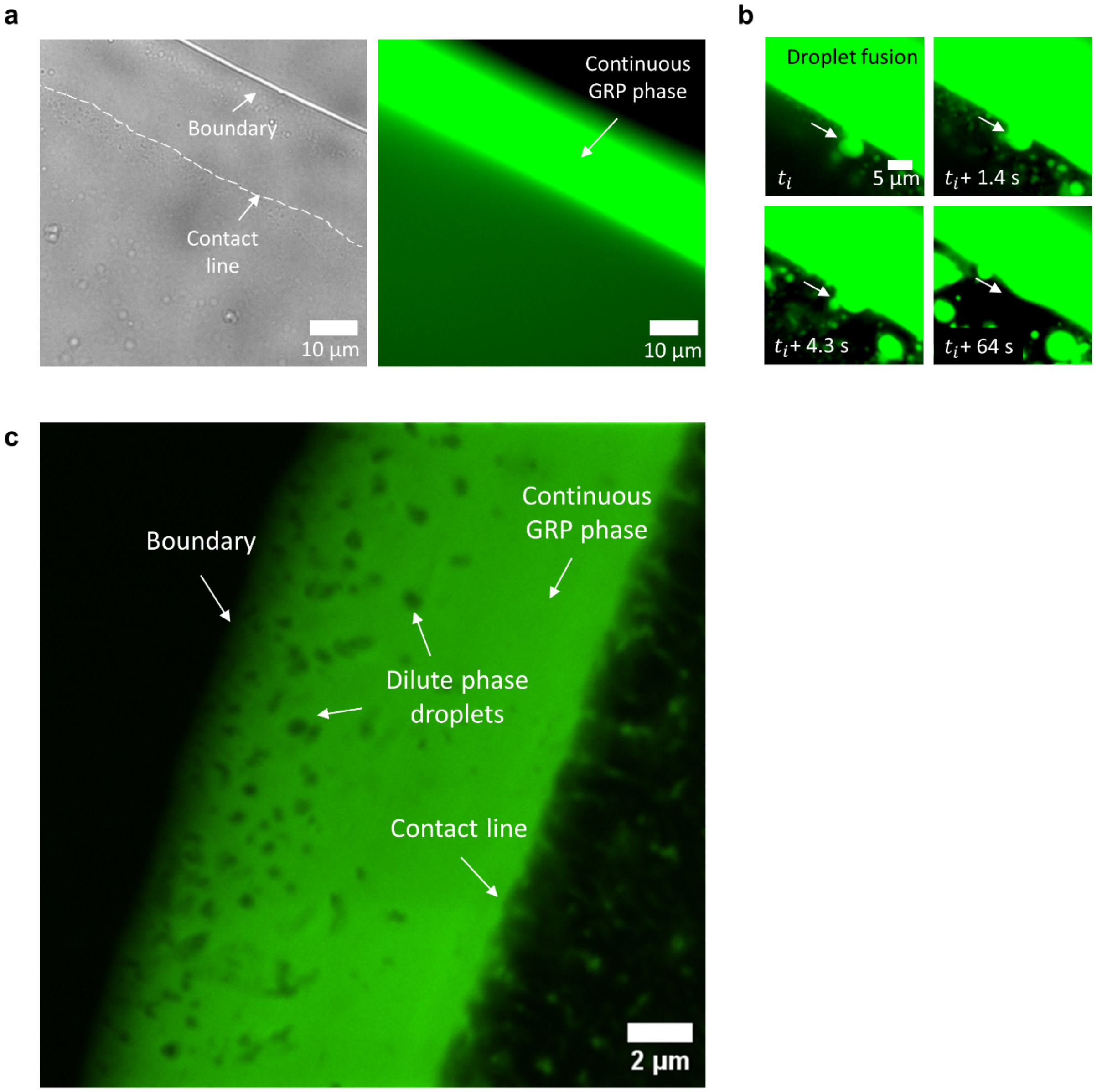


**Supplementary Figure 3. The region between droplet boundary and the contact line is a tick-GRP77-rich condensate phase.** (**a**) Bright-field (left) and corresponding fluorescence image (right) showing the tick-GRP77-rich nature of this region (32 μM starting concentration). (**b**) An example of tick-GRP77 condensate fusing with the continuous tick-GRP77-rich phase. (**c**) Confocal microscopy showing tick-GRP77-depleted aqueous droplets within the inverted phase formed by evaporation assay (125 μM starting concentration).

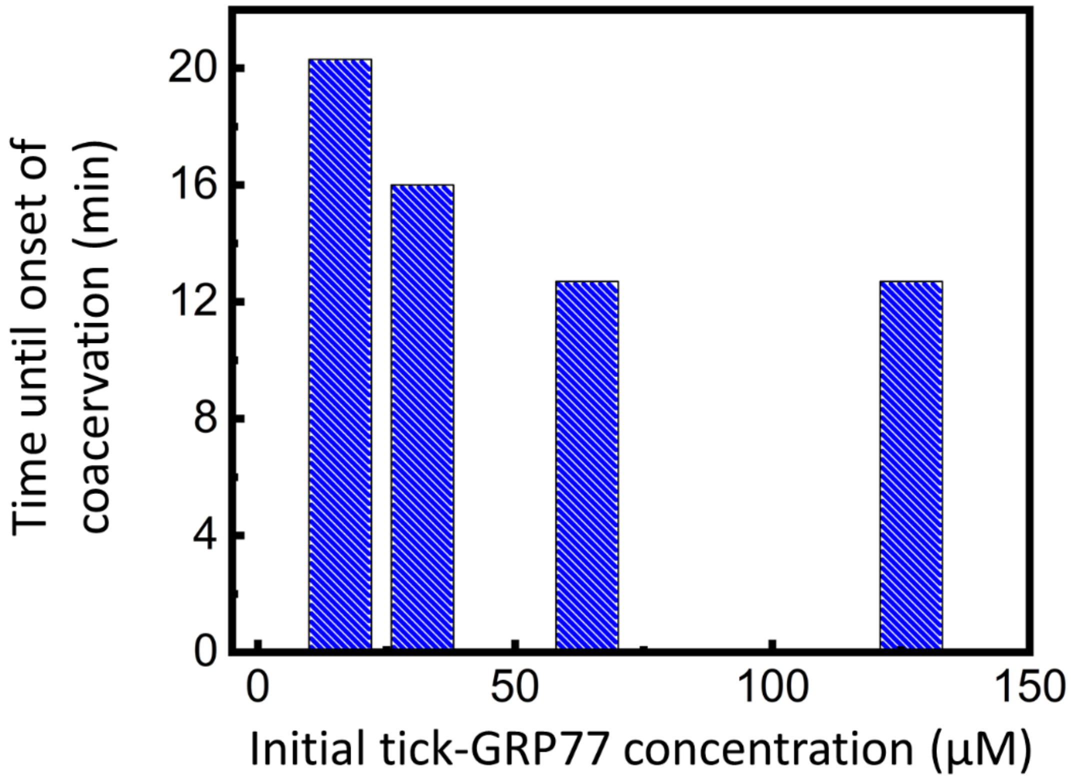


**Supplementary Figure 4: Onset of coacervation is influenced by the starting tick-GRP77 concentration.** The time required for the onset of coacervation is steadily reduced as the initial tick-GRP77 concentration is increased. The values were recorded during droplet evaporation assay in PBS (pH 7.4).

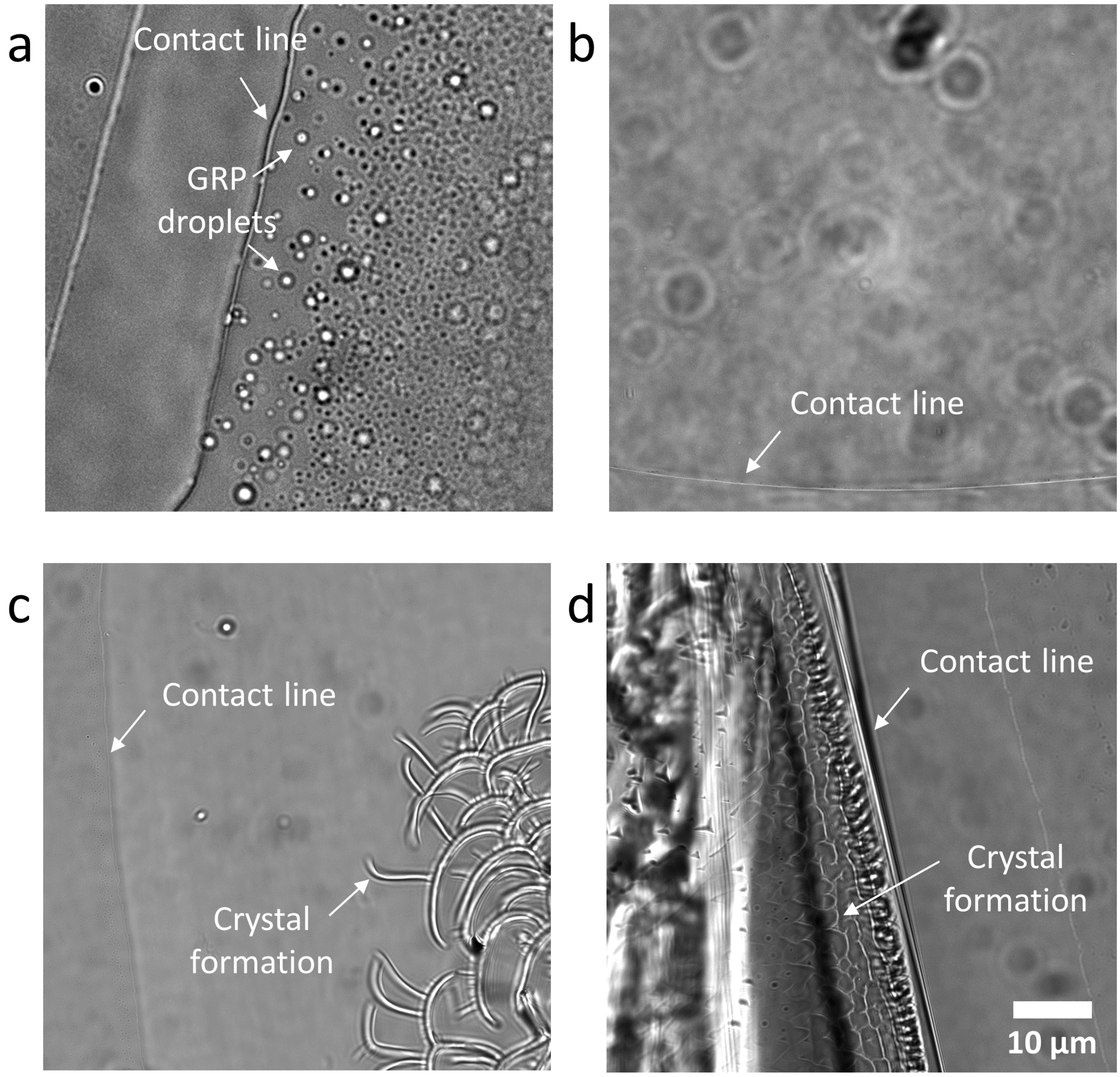


**Supplementary Figure 5: The formed condensates are specific to tick-GRP77 and need salts for their formation.** (**a**) Evaporation of tick-GRP77 (32 µM) in presence of 140 mM NaCl resulted in the formation of condensates. (**b**) On the contrary, evaporation of tick-GRP77 (32 µM) dissolved in pure water did not lead to phase separation. **(c)** Evaporation of globular protein solution, bovine serum albumin (127 µM in PBS, pH 7.4) eventually led to the formation of salt crystals without any phase separation. (**d**) Similarly, just a PBS solution (pH 7.4) did not form condensates but salt crystals.

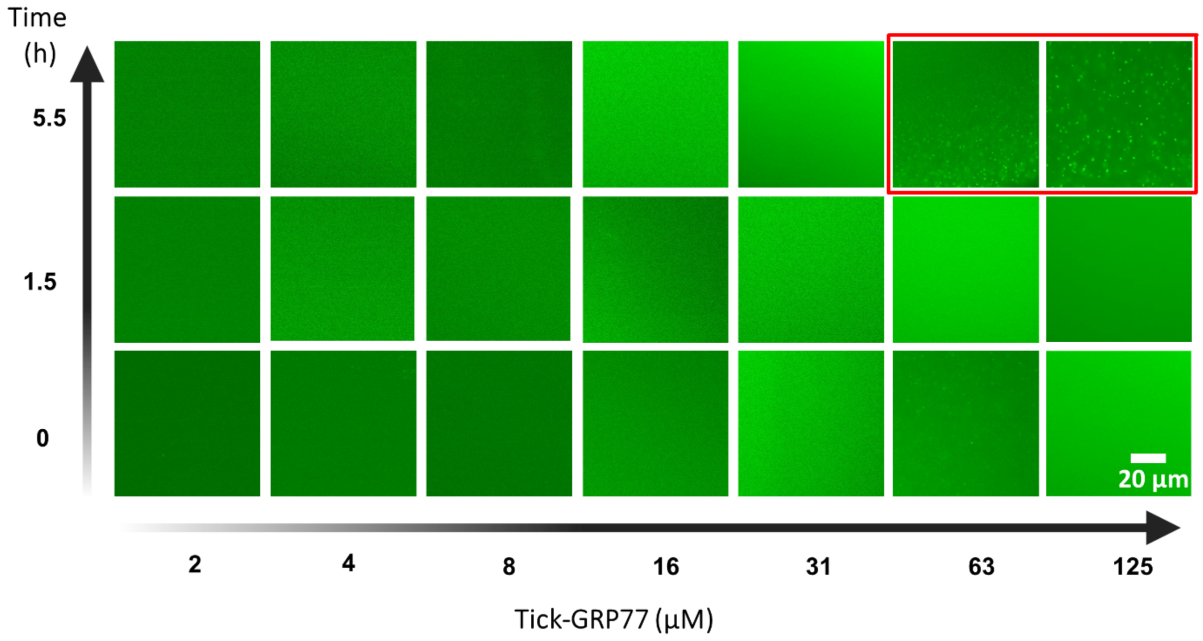


**Supplementary Figure 6. Tick-GRP77 coacervation phase diagram in presence of 0.5 M $Na_2HPO_4$ solution.** Incubation of tick-GRP77 (2–125 µM) at 0.5 M $Na_2HPO_4$ did not lead to instant condensate formation. For concentrations above 63 µM, condensates formed after 5.5 hour of incubation.

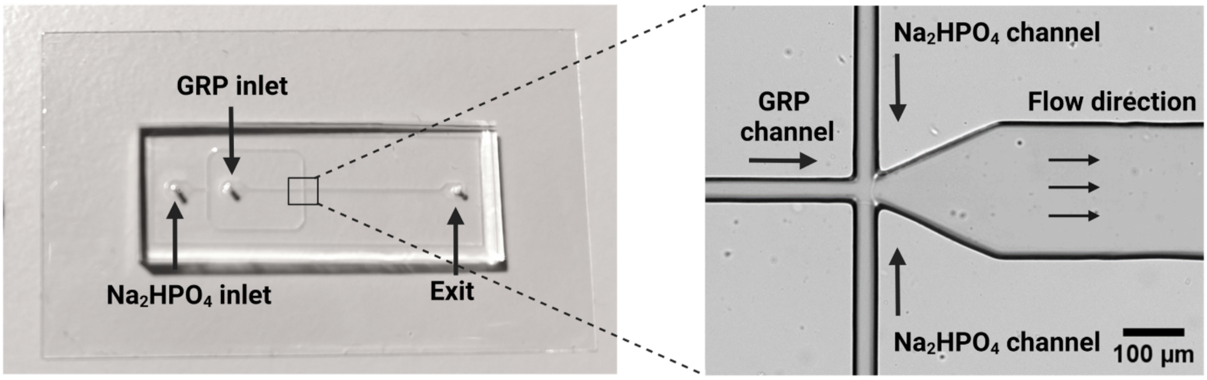


**Supplementary Figure 7: Microfluidic device used in flow-focusing experiments.** Bright-field images showing PDMS-based lab-on-a-chip device with a zoom-in showing the flow-focusing junction.

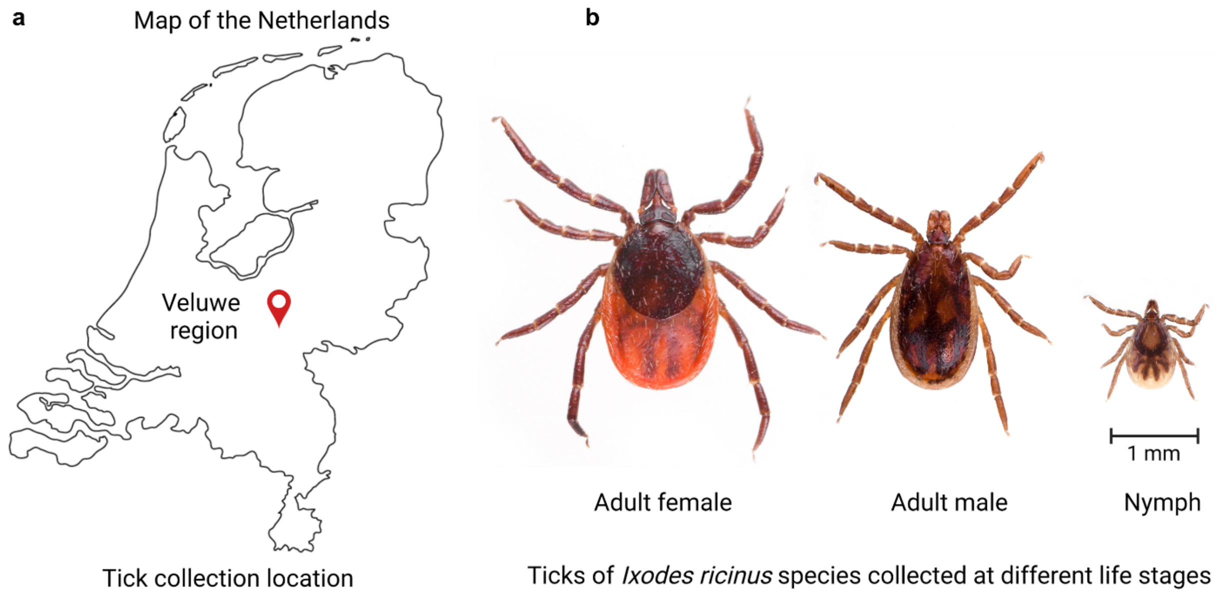


**Supplementary Figure 8. Tick collection region.** (**a**) Location of tick collection in the Netherlands. (**b**) Ticks of species *Ixodes ricinus* at different life stages were collected.

**Supplementary Movie legends**

**Supplementary Movie 1**

Evaporation of a 2 µL sessile droplet of 32 µM tick-GRP77 (doped with 5 mol% OG488-labelled GRP) in PBS (pH 7.4) on a glass slide. The coffee ring effect gradually increases the protein concentration at the droplet boundary inducing the formation of tick-GRP77 condensates.

**Supplementary Movie 2**

Coalescence of two tick-GRP77 condensates. Condensates were obtained by evaporating a 2 µL sessile droplet of 32 µM tick-GRP77 (doped with 5 mol% OG488-labelled GRP) in PBS (pH 7.4) on a PVA-coated glass slide.

**Supplementary Movie 3**

Flow-focusing a stream of 63 µM of tick-GRP77 (doped with 5 mol% OG488- GRP) with two co-flowing 2 M $Na_2HPO_4$ streams leads to the formation of tick-GRP77 condensates at the interfaces of protein-salt streams. The condensates get deformed due to the flow-induced shear.

**Supplementary Movie 4**

Evaporation of a 2 µL sessile droplet with high concentration of tick-GRP77 (500 µM) in PBS (pH 7.4) on a hydrophobic glass slide led to the formation of an interconnected viscoelastic network of condensates near the contact line.

**Supplementary Movie 5**

Time-lapse showing the fluorescent recovery after photobleaching of a fresh (0.5 h old) tick-GRP77 condensate. A small region inside the condensate (obtained by exposing 125 µM tick-GRP77 to 1 M $Na_2HPO_4$ solution) was bleached and showed about 50% recovery of fluorescence intensity within a time span of 5 minutes.

**Supplementary Movie 6**

Time-lapse showing the fluorescent recovery after photobleaching of an aged (18 h old) tick-GRP77 condensate. A small region inside the condensate (obtained by exposing 125 µM tick-GRP77 to 1 M $Na_2HPO_4$ solution) was bleached and showed virtually no recovery of fluorescence intensity within a time span of 5 minutes.
